# Endothelin receptor Aa regulates proliferation and differentiation of Erb-dependant pigment progenitors in zebrafish

**DOI:** 10.1101/308221

**Authors:** Karen Camargo-Sosa, Sarah Colanesi, Jeanette Müller, Stefan Schulte-Merker, Derek Stemple, E. Elizabeth Patton, Robert N. Kelsh

## Abstract

Skin pigment patterns are important, being under strong selection for multiple roles including camouflage and UV protection. Pigment cells underlying these patterns form from adult pigment stem cells (APSCs). In zebrafish, APSCs derive from embryonic neural crest cells, but sit dormant until activated to produce pigment cells during metamorphosis. The APSCs are set-aside in an ErbB signaling dependent manner, but the mechanism maintaining quiescence until metamorphosis remains unknown. Mutants for a pigment pattern gene, *parade*, exhibit ectopic pigment cells localised to the ventral trunk. We show that *parade* encodes Endothelin receptor Aa, expressed in the blood vessels. Using chemical genetics, coupled with analysis of cell fate studies, we show that the ectopic pigment cells derive from APSCs. We propose that a novel population of APSCs exists in association with medial blood vessels, and that their quiescence is dependent upon Endothelin-dependent factors expressed by the blood vessels.

**Lay Abstract:** Pigment patterns are crucial for the many aspects of animal biology, for example, providing camouflage, enabling mate selection and protecting against UV irradiation. These patterns are generated by one or more pigment cell-types, localised in the skin, but derived from specialised stem cells (adult pigment stem cells, APSCs). In mammals, such as humans, but also in birds and fish, these APSCs derive from a transient population of multipotent progenitor cells, the neural crest. Formation of the adult pigment pattern is perhaps best studied in the zebrafish, where the adult pigment pattern is formed during a metamorphosis beginning around 21 days of development. The APSCs are set-aside in the embryo around 1 day of development, but then remain inactive until that metamorphosis, when they become activated to produce the adult pigment cells. We know something of how the cells are set-aside, but what signals maintain them in an inactive state is a mystery. Here we study a zebrafish mutant, called *parade*, which shows ectopic pigment cells in the embryo. We clone the *parade* gene, identifying it as *ednraa* encoding a component of a cell-cell communication process, which is expressed in blood vessels. By characterising the changes in the neural crest and in the pigment cells formed, and by combining this with an innovative assay identifying drugs that prevent the ectopic cells from forming, we deduce that the ectopic cells in the larva derive from precocious activation of APSCs to form pigment cells. We propose that a novel population of APSCs are associated with the blood vessels, that these are held in a quiescent state by signals coming from these vessels, and that these signals depend upon *ednraa*. Together this opens up an exciting opportunity to identify the signals maintaining APSC quiescence in zebrafish.

## Introduction

Pattern formation is a crucial aspect of development since it creates the functional arrangements of cell-types that allow an organism to thrive. Pigment pattern formation – the generation of correctly distributed pigments or pigmented cells within the skin or elsewhere in the body – is a case in point, with pigmentation crucial for diverse aspects of an animal’s ecology, including avoidance of predators, kin recognition, mate selection, thermal regulation and UV protection.

In vertebrates, all pigment cells except those of the pigmented retinal epithelium, are derived from a transient embryonic tissue called the neural crest. Neural crest cells are multipotent, generating numerous types of neurons, glia, pigment cells and other derivatives. They are also highly migratory, moving from their origin in the dorsal neutral tube to occupy diverse sites throughout the embryo. Thus, correct positioning of the different cell-types is a crucial aspect of their development.

Pigment cells in mammals consist only of melanocytes, making (and secreting) black eumelanin or yellow pheomelanin granules. In fish, amphibians and reptiles, pigment cells are much more diverse [1], allowing the generation of the varied and often beautiful pigment patterns these groups display. The zebrafish *Danio rerio* has rapidly become a paradigmatic example for the genetic and cellular study of pigment pattern formation [2–6]. Zebrafish pigment patterns consist of three pigment cells, black melanocytes making melanin, yellow xanthophores making pteridines and carotenoids, and iridescent iridophores containing reflecting platelets [1].

Zebrafish, in common with most fish, develop two distinct pigment patterns, an early larval pigment pattern generated in the embryo by direct development of pigment cells from neural crest cells, and an adult pattern formed during metamorphosis, mostly through the de novo differentiation of pigment cells from adult pigment stem cells (APSCs, also formerly known as melanocyte stem cells; [7–9]). The adult pigment pattern consists of prominent stripes consisting of melanocytes and associated blue iridophores, alternating with pale stripes (interstripes) consisting of dense silver iridophores and xanthophores. Pigment pattern formation in adults is partially well characterised, with many genes identified that regulate the production of different pigment cell-types from the APSCs or which control their cellular interactions to create the bold horizontal stripe pattern. A key aspect of adult pigment pattern formation that is less well-understood is the generation of the APSCs from neural crest cells. Remarkably, elegant experimental studies from multiple laboratories have established that these are set-aside from the neural crest in a narrow time-window (9-48 hours post-fertilisation (hpf); [10]), with at least some occupying a niche within the dorsal root ganglia (DRGs; [9, 11]) of the peripheral nervous system. They remain quiescent until metamorphosis begins around 20 days post-fertilisation (dpf), when they become activated. The mechanisms controlling their quiescence and their activation are largely unknown.

Pigment pattern formation in embryos is also poorly understood[3]. The embryonic pigment pattern consists principally of four longitudinal stripes of melanocytes, with iridophores arranged in a characteristic association with the melanocytes in three of these (Dorsal, Ventral and Yolk Sac Stripes), whereas the Lateral Stripe consists only of melanocytes; xanthophores then occupy the space under the epidermis between these stripes. Whereas in the adult the stripes are in the dermis, in the embryo they are associated with other structures, including the CNS, horizontal myoseptum of the body muscle blocks, the internal organs and the ventral most yolk sac. One study has investigated the detailed mechanism driving the association of melanocytes with the horizontal myoseptum [12]. In general, stripes form through migration of pigment cell precursors down migration pathways used by neural crest. These neural crest migration pathways are known as the dorsal (or dorsolateral) migration pathway, consisting of cells migrating under the epidermis and over the outer face of the somites/developing muscle blocks, and the medial migration pathway, running between the neural tube and notochord and the medial face of the somites/developing muscle blocks [13]. Pigment cell precursors of different fates use distinct migration pathways. Thus, xanthoblasts only use the lateral migration pathway, iridoblasts use only the medial migration pathway, whereas melanoblasts use both [14–17]. Note that migrating pigment cell precursors often show early signs of pigmentation i.e. they are differentiating as they migrate. During the migration phase, early differentiating melanocytes and iridophores can be found in the ventral trunk on the medial migration pathway, but these have disappeared by 72 hpf as those cells migrate into the Ventral and Yolk Sac Stripes[16].

As in adult pigment pattern formation, mutants affecting embryonic pigment pattern offer an exciting entry-point to the study of the mechanisms controlling pigment pattern formation. In a large-scale ENU mutagenesis screen performed in 1996, we identified two zebrafish mutant alleles, *pde^tj262^* and *pde^tv212^*, that defined the *parade (pde)* gene [18]. These mutants showed ectopic melanophores and iridophores in a well-defined region of the ventral side of the posterior trunk (Fig.1B and E), in addition to a stripe pattern similar to the wild type (WT) pigment phenotype (Fig. 1A and C). The striking coincidence of melanin and reflecting platelet distribution lead us to initially propose that the *pde* mutant phenotype results from differentiation of pigment cells of mixed fate i.e. with both melanin granules and reflecting platelets within the same cell [18]. Here we perform a comprehensive analysis of the *pde* mutant phenotype. We show that the *pde* locus encodes a zebrafish Endothelin receptor A, Ednraa, which is expressed only in the developing blood vessels. Using chemical genetics, coupled with analysis of cell fate studies, we propose that APSCs occupy a niche associated with the medial blood vessels of the trunk, likely to be a second PNS niche within the sympathetic ganglion chain, and that these become precociously activated in *pde/ednraa* mutants. Thus we hypothesise that key components of that blood-vessel niche are Ednraa-dependent factors that promote APSC quiescence.

**Figure 1.**
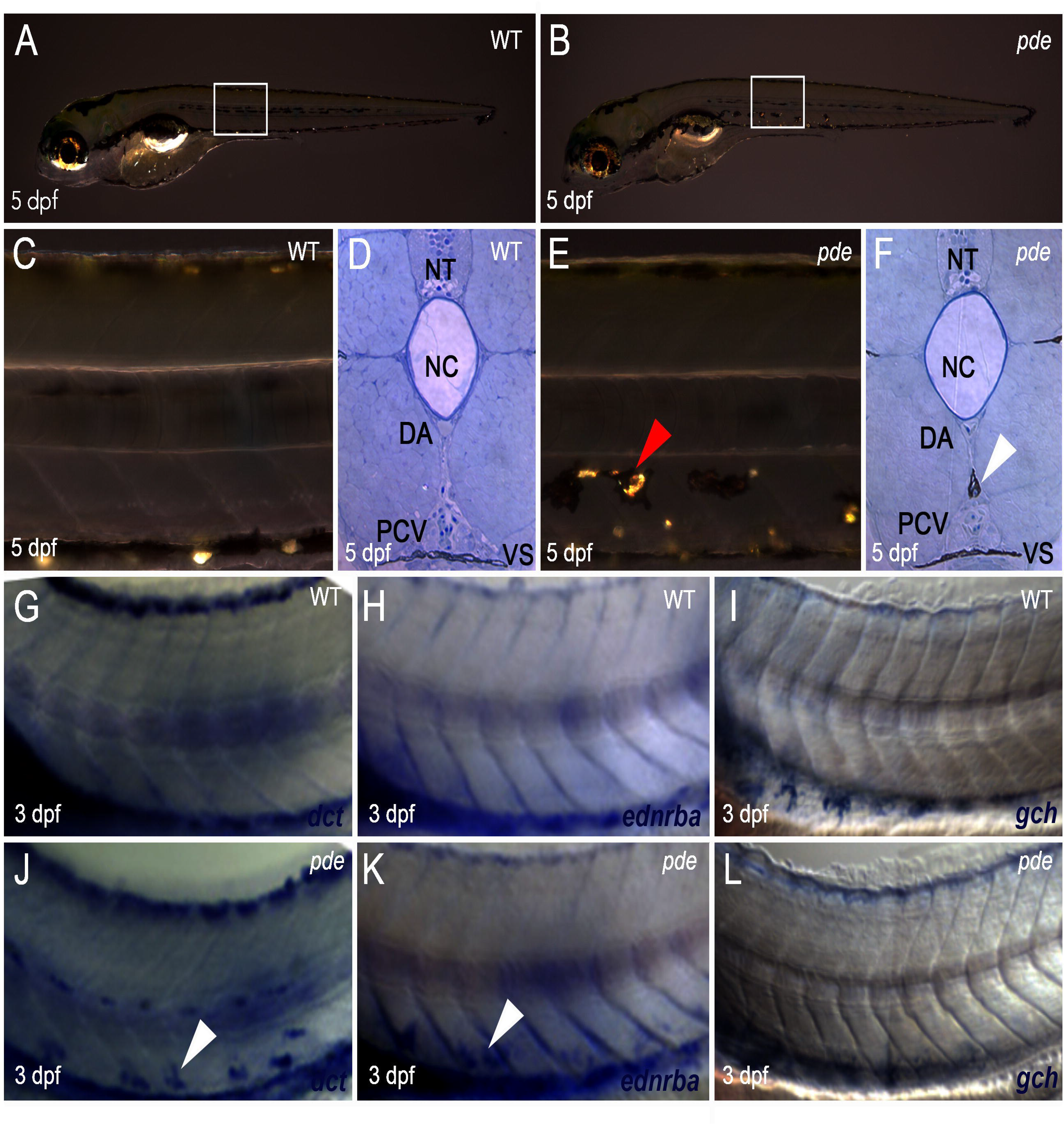
*pde* mutants display ectopic melanocytes and iridophores, but not xanthophores, in the ventral medial pathway. (A,B) Overview of early larval WT (A) and *pde* (B) pigment phenotype at 5 dpf. (C-F) Anatomical location of ectopic pigment cells in *pde*. Magnification of lateral views (white boxes in A and B) and cross sections of posterior trunks show no pigment cells on the medial migration pathway in the ventral trunk of WT larvae (C and D). Ectopic pigment cells are located in the ventral trunk of *pde* mutants (E; red arrowhead), under the dorsal aorta (DA) and above the posterior cardinal vein (PCV) as shown by cross sections (F, white arrowhead). (G-L) Whole mount in situ hybridization of 3 dpf WT (G-H) and *pde* mutants (J-L) embryos for *dct* (G and J), ednrba (H and K), and *gch* (I and L). Ectopic *dct* (J; white arrowhead) and *ednrba* (K; white arrowhead) expression is seen in the ventral trunk of *pde* mutant larvae. Neural tube (NT), notochord (NC) and ventral stripe (VS).

## Materials and Methods

### Fish husbandry

Fish care and procedures were approved by the University of Bath Ethical Review Committee, and were performed in compliance with the Animals Scientific Procedures Act 1986 of the UK. WT strain AB, *pde^tj262^, pde^tv212^* and *pde^hu4140^, Tg(fli1a:EGFP)^y1^* and *Tg(-4725sox10:cre)^ba74^*; *Tg(hsp:loxp-dsRed-loxp-EGFP)* were used.

### Genetic mapping

#### Mapping panels

Two specific sets of microsatellite markers, the G4 and H2 panels (Geisler et al., 2007) were used for bulk segregant analysis of *parade*, placing the mutation on linkage group 1. We subsequently used a consolidated meiotic map of the zebrafish genome, ZMAP, which is available on ZFIN (http://zfin.org/cgi-bin/webdriver?MIval=aa-crossview.apg&OID=ZDB-REFCROSS-010114-1; Sprague et al., 2006; Geisler et al., 2007).

#### Reference zebrafish line

The standard mapping wild-type line WIK was crossed with mutant *parade^tj262^* carriers (in AB wild-type background). This founder generation F0 was incrossed to produce heterozygous F1. Eight pairs of F1 fish were incrossed to produce the F2 generation. In total 1796 F2 embryos at 5 dpf were sorted into two groups, those that display the mutant phenotype and those with a normal wild-type pigment phenotype. These were transferred into 96-well plates where the extraction of genomic DNA was performed and stored. Additionally, genomic extracts of 10 homozygous *parade^tj262^* and 10 wild-type sibling embryos of each of the 8 parent pairs were pooled for identification of new SSLP markers.

#### Mapping procedure

The *pde* locus was mapped to linkage group 1 (LN 1) by bulked segregant analysis using pooled genomic DNA of 48 wild-type siblings and 48 *parade^tj262^* mutant embryos of the F2 generation obtained from F1 Pair 2. For fine mapping we designed new mapping primers using the Primer3 online software tool (http://frodo.wi.mit.edu/primer3/, default settings plus 1 primer pair per 300-400 bp, (Rozen and Skaletsky, 2000)). Primer pairs were designed to generate PCR products of 300–400 bp, which if generating polymorphic PCR amplicons, were then tested on the 1796 single F2 embryos to determine the frequency of recombination.

#### PCR protocol

2 μl of individual or pooled DNA were added to 13 μl of PCR mix (1.5 μl 10 x buffer for KOD Hot Start DNA Polymerase; 1.5 μl dNTP’s with 0.2 mM for each nucleotide; 1 μl primer mix with 10 mM forward and 10 mM reverse primer; 0.3 μl DMSO; 0.6 μl 25 mM MgSO4; 8.3 μl MiliQ water; 0.3 μl KOD polymerase (Novagen)). PCR’s were run in a GStorm thermal cycler (Gene Technologies Ltd) (program: 2 min 94° C; then 35 x (30 sec 94° C, 30 sec 60° C, 30 sec 72° C); then 5 min 72° C; then stored at 10° C). The PCR products were then analysed by electrophoresis.

### Morpholino injection

Custom morpholinos were purchased from Gene Tools LLC (Philomath, USA) and resuspended in autoclaved MilliQ water to a stock dilution of 20 g/ l. Stock solutions were stored at −20° C to prevent evaporation and heated at 70° C for 5 min prior to dilution to eliminate precipitates. Wild-type embryos were injected into the yolk at the 1-cell stage with 5 ng/nl dilutions. Phenotypes were observed under the microscope at 3 dpf and 4 dpf. Volumes varied between 5 and 10 nl per embryo. Sequences of morpholino oligos can be found in Table 1.

**Table 1.**
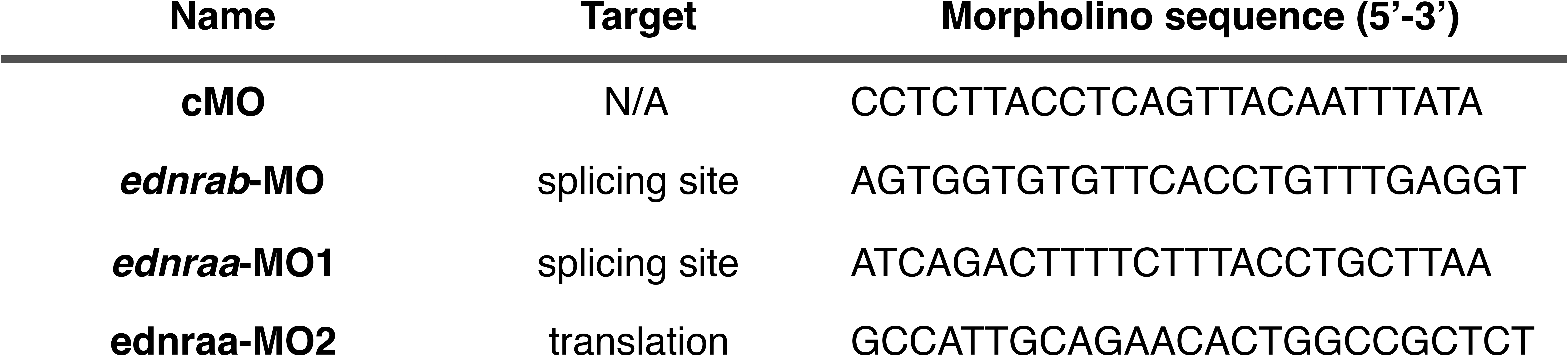
Morpholino sequences. A random sequence morpholino (cMO) provided by Gen Tools and a splice-blocking morpholino against *ednrab* (*ednrab*-MO) were used as control. *ednraa* was targeted with either a splice-blocking morpholino (*ednraa* -MO1) or a translation-blocking *ednraa* morpholino (*ednraa*-MO2).

### Whole mount *in situ* hybridization

Our protocol for whole-mount in situ hybridization is based on (Thisse et al., 1993). All solutions were prepared with DEPC-treated autoclaved MiliQ water or PBS. All probes for in situ hybridization were synthesized using the Dig RNA Labeling Kit (Boehringer). Depending on the orientation of the gene and choice of plasmid, we have chosen either T3, T4 or SP6 RNA polymerases for creating the antisense RNA probe.

Briefly, embryos between 5 and 120 hpf were euthanized with a lethal overdose of tricaine and fixed overnight (4 % PFA in PBS). Embryos older than 18 hpf were dechorionated with forceps before fixation, younger embryos were dechorionated with flamed forceps after fixation. All following steps were carried out in 1.5 ml microfuge tubes with 1 ml liquid volumes; up to 40 embryos were processed per tube. The fixative was rinsed out with PBTween (DEPC-treated PBS plus 0.1 % Tween). Embryos were then gradually dehydrated on ice with 25, 50, 75 and 100 % methanol. For the ISH procedure, embryos were rehydrated on ice using 5 min washes with 75, 50, 25 % methanol followed by 3 × 5 min washes with PBTween at room temperature. To improve permeability, embryos between 30 – 48 hpf were treated for 10 min with 0.01 mg/ml proteinase K/PBTween, embryos; older than 50 hpf were treated for up to 20 min. The proteinase K was removed by washing samples for 3 × 5 min with PBTween followed by a brief re-fixation step with 4% PFA for 20 min at room temperature. The fixative was then washed out with 3 × 5 min PBTween. To prepare the samples for optimal binding conditions, embryos were prehybridized with hybridization mix (formamide, 20 × SSC, heparine, tRNA, Tween20, citric acid; stored at −20° C) in a water bath at 65° C for 4 h. The hybridization of the RNA probe (diluted 1:400) was done over night at 65° C in 200 μl hybridization mix. The samples were washed with decreasing concentrations of hybridization mix (100 % hyb mix; 75 % hyb mix/25 % 2x SSCT; 50 % hyb mix/50 % 2x SSCT; 25 % hyb mix/75 % 2x SSCT; 100 % 2x SSCT), each for 10 min, followed by 2 × 30 min washing with 0.2x SSCT. All steps were carried out in a water bath at 65° C. Samples were then infiltrated for 3 × 5 min with MABT (from 5 x MAB stock solution plus 0.1 % Tween) at room temperature. Samples were blocked for 3 h in MABTween with 5 % sheep serum at room temperature under gentle shaking. Antibody binding reaction was performed over night at 4° C using 200 μl of 1:5000 diluted anti - dioxigenin alkaline phosphatase conjugated antiserum in blocking solution. Samples were then washed for 6 × 15 min in MABT, then prepared for signal detection reaction by infiltrating for 3 × 5 min with NBT/BCIP buffer (100 mM Tris HCl pH 9.5, 50 mM MgCl2, 100 mM NaCl, 0.1 % Tween). The embryos were transferred into 9-well glass dishes to observe the staining reaction under the dissecting microscope. 300 μl of BMPurple substrate solution were added per well and the development of the reaction regularly monitored. Signal development was stopped by washing with PBTween.

### Chemical screening

#### Compounds used

The compounds used in this investigation originated from the 1280 compounds from the LOPAC library (Sigma L01280), 80 compounds from a kinase inhibitor library (Biomol 2832) and 33 compounds from a phosphatase inhibitor library (Biomol 2834A). The compound libraries were stored at −80C and plates thawed at room temperature prior to use. For screening potential effects on developing zebrafish embryos compounds were diluted to 10uM in E3 embryo medium.

For drug treatments, six embryos were placed in each well of 24 well plates. Chemical treatments were prepared in 10uM doses in 1ml E3 medium. To ensure that compounds did not precipitate, the plates containing small molecule treatments were placed on The Belly Dancer (Sorval) for a minimum of 20 minutes. When embryos reached 8hpf E3 medium was removed and 1ml of E3 containing 10uM of compound was added to each well. Wells containing E3 only and 10uM DMSO were used in each chemical treatment as controls. Plates containing treated embryos were incubated at 28C until the embryos reached 5dpf. Detailed observations were recorded daily and any dead embryos removed to avoid contamination of the medium. At day 5 embryos were anaesthetised with a small dose of tricaine to enable easy manipulation for accurate detailing of pigment phenotype. Pigment cell phenotypes and rescue of the *parade* mutant phenotype were recorded as well as additional phenotypes such as developmental abnormalities (e.g. shortening of the tail or cardiac edema).

#### Re-screening of important chemical ‘hits’

Once molecules of specific interest to pigment cell development were identified the treatment was repeated on embryos at different developmental time points and across a concentration gradient in order to identify the developmental time point of greatest phenotypic significance and the optimal small molecule concentration.

### Whole mount immunostaining

Antibody staining was largely performed as described in [19]. Prior to primary antibody incubation, embryos were permeabilized with 1 ml of 5 μg/ml of proteinase K/distilled water during 30 minutes at 37 ºC, then rinsed with 5% of goat serum/distilled water at RT for 5 minutes and washed 3×1 hour with distilled water at RT. The antibodies used were Hu (1:500,) and Alexa Fluor 488 goat-mouse IgG (1:750, Molecular Probes, Cat. # A11001).

### Image acquisition and processing

For in vivo microscopy, embryos were mounted at the Leica Fluo III or Zeiss Axiovision dissection microscope in 0.6 % agarose or 3 % methyl cellulose. For anaesthesia, 0.02 % Tricaine was added freshly to the mounting medium. The embryos were transferred to a microscope slide or to a glass-bottomed Petri dish. For immunofluorescence, embryos were visualized with a Nikon Eclipse E800 or Zeise M2 Imager compound microscope under bright field, incident light or epifluorescence illumination, mainly using a 10x, 20x and 63x objective for magnification. Images were taken with a DS-U1 (Nikon) or Orca and color 546 camera (Zeiss). Confocal fluorescence imaging was performed on an inverted Zeiss LSM 510 Meta or LSM 880 confocal microscope with 20x and 40x objectives. Incident light was provided by installing an additional antler lamp (Leica) around the microscope to allow visualisation of iridophores. NIS-Elements D2.30, Zen blue, Fiji and/or Adobe Photoshop 7 were used to adjust white balance and exposure and to assemble the images to figures.

## Results

### Early larval *pde* mutants have supernumerary iridophores and ectopic melanocytes and iridophores

To test our initial working model, that the ectopic cells are cells of mixed melanocyte/iridophore fate, we began by asking exactly where the ectopic cells were located, and which pigment cell-types were involved. Whilst WT siblings at 5 dpf (Fig.1D) show no pigment cells in the ventral trunk between the notochord and the Ventral Stripe, in *pde^tj262^* mutants (from now on referred to simply as *pde* unless specified otherwise) ectopic chromatophores are located in a medial position, directly beneath the dorsal aorta (DA) and above the posterior cardinal vein (PCV; Fig. 1F). Thus, their location corresponds precisely to the ventral region of the medial neural crest migration pathway.

In zebrafish, pigment cell migration (both as unpigmented progenitors and as pigmenting cells) is pathway specific, with cells of the melanocyte and iridophore, but not xanthophore, lineages using the medial pathway [3]. Direct observation clearly shows cells with the pigmentation characteristics of both melanocytes and iridophores in this region of *pde* mutants, but detection of xanthophores by their pigmentation is difficult at these stages. Thus, to assess the presence of ectopic xanthophores, we used whole-mount in situ hybridisation (WISH) using fate-specific markers. Detection of the melanocyte marker *dopachrome tautomerase* (*dct*; Fig. 1J), the iridoblast marker *endothelin receptor ba* (*ednrba*; Fig. 1K) and the xanthophore marker *GTP cyclohydrolase I* (*gch*; Fig. 1L) at 72 hpf, provided clear confirmation that *pde* mutants display ectopic melanocytes and iridophores, but importantly revealed the complete absence of ectopic xanthophores in the ventral trunk (Fig. 1L).

Initial observations of *pde* mutants indicated the intriguing possibility that some of the ectopic chromatophores displayed mixed characteristics of both iridophores and melanophores (Fig. 1E and F; [18]). This suggested that the phenotype might result, at least in part, from a failure of the normal mutual repression of alternative pigment cell fates. To test this, we used transmission electron microscopy to assess the structure of the ectopic cells. Unexpectedly, we consistently saw that melanosomes, the melanin synthesising organelles of melanocytes, and the reflecting platelets that contain the reflective crystals in iridophores, formed separate clusters, and were never intermingled (Fig. 2A-2C). Furthermore, these clusters were consistently separated from each other by double membranes (Fig. 2A-2C), similar to the appearance of melanocytes and iridophores in the WT Yolk Sac Stripe (Supp. Fig. S1).We conclude that the ectopic pigment cells are not of mixed fate, but instead are tightly associated individual melanocytes and iridophores; we note that this arrangement is characteristic of iridophores in their normal locations, where in each of the Dorsal, Ventral and Yolk Sac Stripes iridophores are tightly associated with melanocytes.

**Figure 2.**
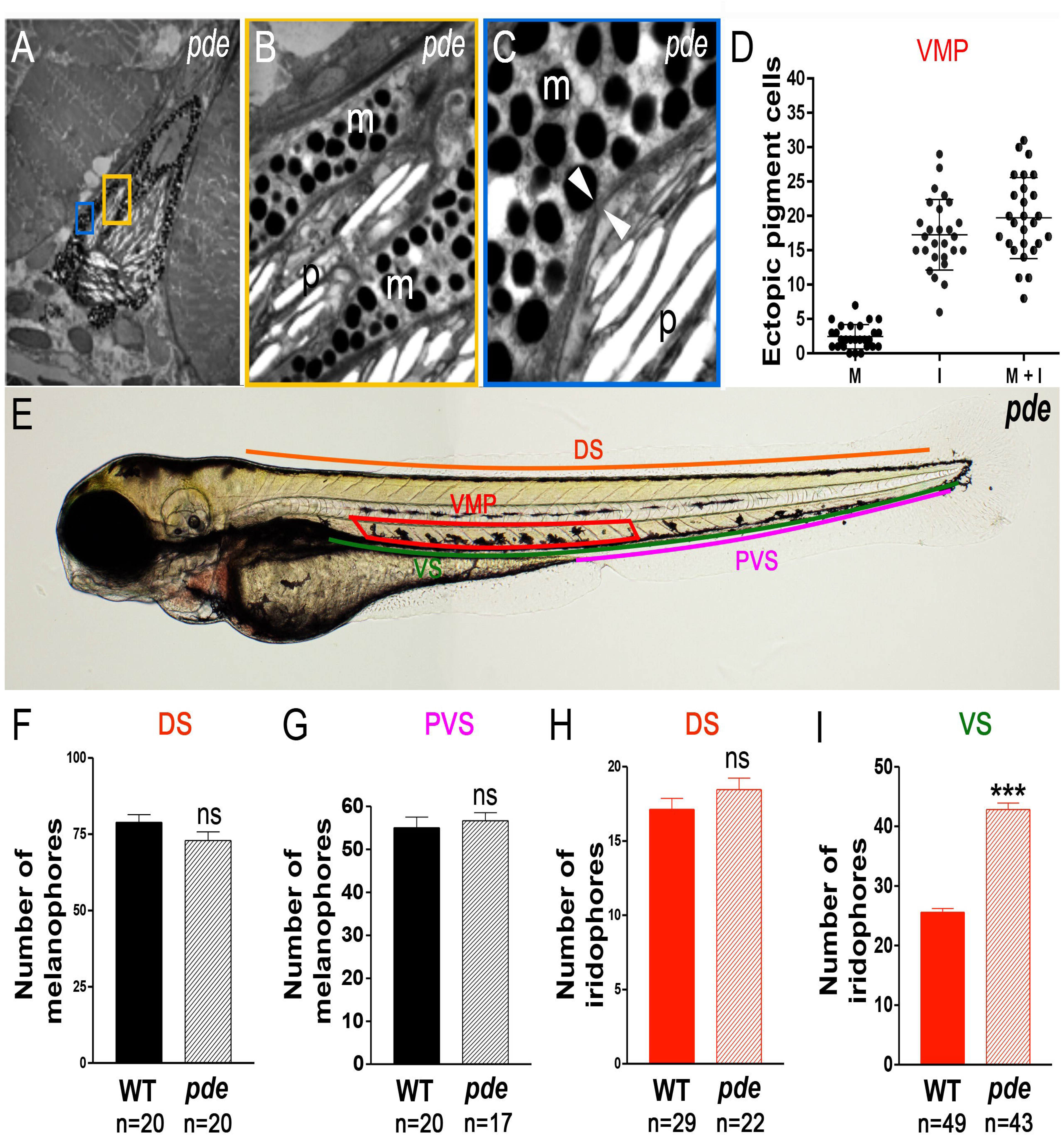
*pde* mutants show supernumerary melanocytes and iridophores in the Ventral Stripe and nearby medial migration pathway, but not the Dorsal Stripe. (A-C) Transmission electron photomicrographs of ectopic pigment cells in *pde* mutants. Magnifications of yellow (B) and blue (C) boxes show melanosomes (m) and reflecting platelets (p) separated by a double membrane (C; white arrowheads). (D) Dot-plot of quantitation of ectopic melanocytes (M), iridophores (I) and overall number of ectopic pigment cells (M+I) reveals a variable phenotype, with a consistently larger number of iridophores than melanocytes in the ectopic position (iridophores mean + s.e.=17.26±0.99; melanocytes mean + s.e.=2.44±0.34, n=27). (E) Regions where number of pigment cells were counted, Dorsal Stripe (DS; orange line), Ventral Stripe (V; green line), posterior Ventral Stripe (PVS; pink line) and ventral medial pathway (VMP, red box). (F-I) Quantitation of number of melanocytes in dorsal (F; p>0.05, two-tailed t-test, WT mean + s.e.= 78.8+2.5, n=20 and *pde* m + s.e.=72.8 + 2.8, n=20) and posterior ventral stripes (G; wild-type: mean + s.e. 55.0 + 2.5 n=20; *pde*; 56.6 + 1.8, n=17) show no significant (ns) difference between WT and *pde* mutants. Iridophore quantitation in the DS (H; p>0.05, two tailed t-test, WT mean + s.e. = 22.9 + 0.9, n=29; *pde* mean + s.e. = 20.4 + 1.1, n=22) is not different between WT and *pde* mutants while the ventral stripe has a 58% increased number of iridophores compared to WT embryos (I;***, p<0.0001, two-tailed t-test; WT mean + s.e. = 25.5 + 0.6, n=49; *pde* mean + s.e. = 42.8 + 1.0, n=43).

We hypothesised that the ectopic cells might reflect a defect in pigment cell migration through the ventral pathway, with cells that would normally contribute to the Ventral Stripe becoming stuck during migration. To test this, we quantitated the ectopic pigment cells and the cells of both the Dorsal and Ventral Stripes (Fig. 2D-I). Our model predicted that the counts in the Dorsal Stripe would be unaffected, but that pigment cells in the Ventral Stripe might be reduced, perhaps in proportion to the number of ectopic cells. Quantitation of the ectopic pigment cells (melanocytes + iridophores) in the ventral medial pathway revealed a variable phenotype, with a consistently larger number of iridophores than melanocytes in the ectopic position (Fig. 2D). The number of melanocytes in the Dorsal (Fig. 2F) and Ventral (Fig. 2G) Stripes in *pde* mutants was not statistically different to those of WT siblings. Similarly, there was no significant difference in the number of iridophores in the DS of *pde* mutants and WT siblings (Fig. 2H). Interestingly, and in contradiction to the disrupted migration model, the number of iridophores in the ventral stripe of *pde* mutants shows a 58% *increase* compared to WT siblings (Fig. 2I). Thus, we reject the disrupted migration hypothesis, and instead note that *pde* mutants display a regionally localised increase in melanocytes and iridophores in the ventral trunk, with some as supernumerary cells in the Ventral Stripe, but many in an ectopic position nearby.

### *pde* encodes the endothelin receptor Aa gene

To identify the mutation that causes the *pde* phenotype we crossed *pde* heterozygotes onto a WIK background, and mapped the locus of the mutation using two sets of microsatellite markers, G4 and H2 [20]. The *pde* mutation showed strong linkage to markers Z15424 and Z23059 on chromosome 1, ~1.5 cM and ~3.8 cM away from the mutation, respectively (Fig. 3A). Using the 8^th^ version of the zebrafish genome assembly, further analysis showed that the marker P249 in clone BX51149 was only 0.2 cM (4 recombinants in 17959 embryos) away from the mutation, placing the *pde* mutation about 132 kb away in clone CU462997, in which three genes were annotated: 1) *mineralocorticoid receptor* (now renamed *nuclear receptor subfamily 3, group C, member 2, nr3c2*), 2) *Rho GTPase activating protein 10* (*arhgap10*), and 3) *endothelin receptor Aa* (*ednraa*). Of these three candidate genes, *ednraa* has been previously reported to be required in the development and patterning of the neural crest-derived ventral cranial cartilages. Furthermore, *ednraa* is expressed in the developing blood vessels of the posterior trunk [21]. This striking correlation between *ednraa* expression and the region with ectopic pigment cells in *pde* mutants made *ednraa* a strong candidate. Previous studies had not reported a pigment phenotype during morpholino-mediated knockdown, but the focus in that work had been on craniofacial development, reflecting another region of *ednraa* expression [21].

**Figure 3.**
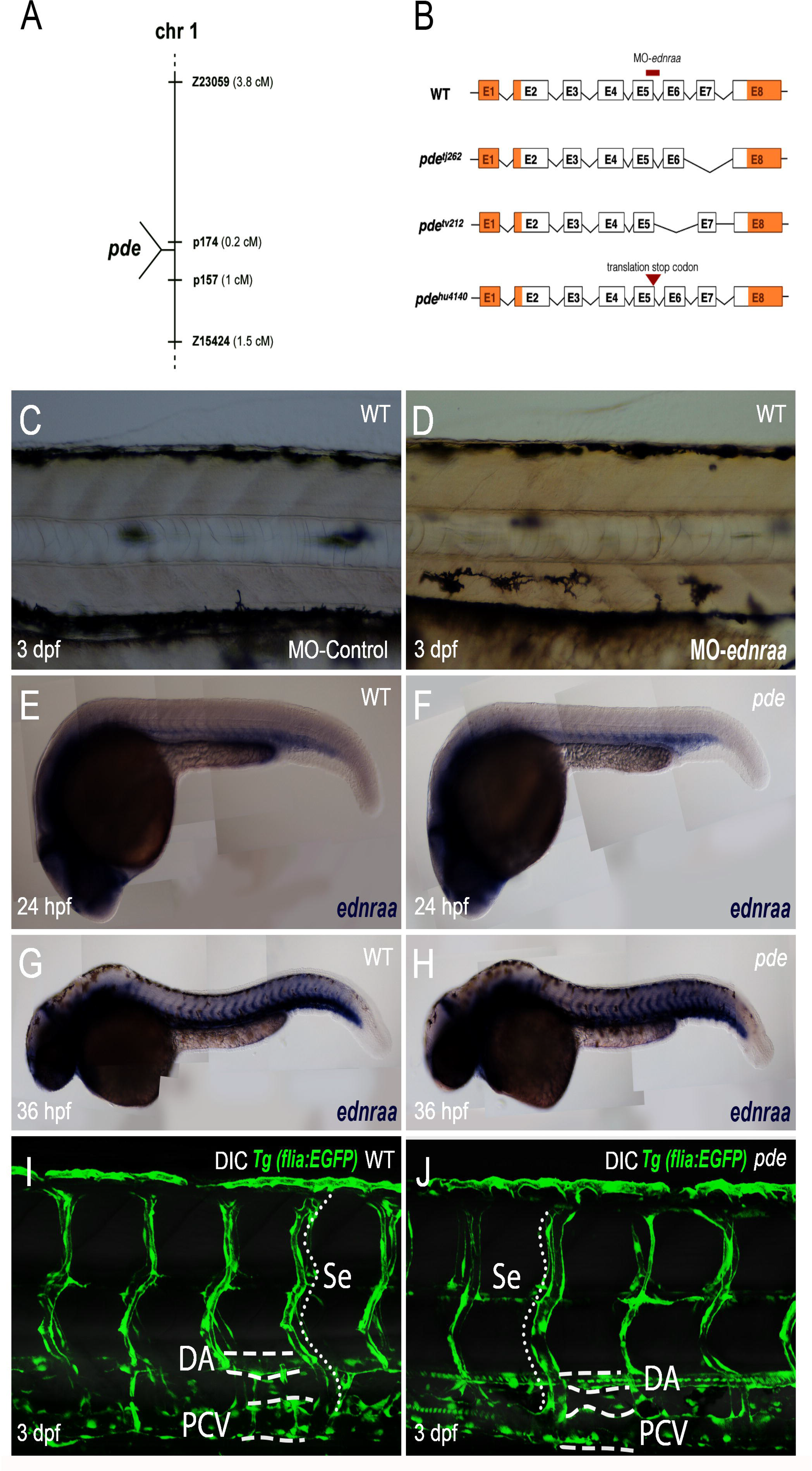
*pde* mutations affect *ednraa*, but not blood vessel formation and patterning. (A) *pde* map position on chromosome 1. (B) Schematic of predicted mRNA structure based upon sequencing of cDNA from *pde^tj262^, pde^tv212^* and *pde^hu4140^* mutants. cDNA of *pde^tj262^* mutants lack exon 7, *pde^tv212^* lack exon 6 and *pde^hu4140^* have a transition mutation (AGA847TGA) that causes a premature translation stop (triangle) in the 3’ region of exon 5. The location of the *ednraa* splice-blocking morpholino (MO-*ednraa*)is indicated (red bar). (C-D) Injection of *ednraa* morpholino into WT embryos phenocopies *pde* mutant pigment phenotype. (C) WT embryos injected with a control morpholino (MO-control) display a normal phenotype. (D) WT sibling injected with MO-*ednraa* display ectopic pigment cells in the ventral medial pathway. (E-H) Whole mount *in situ* hybridization of *ednraa* at 24 hpf and 36 hpf is restricted to the developing blood vessels, and is indistinguishable between *pde* mutants (F and H) and their WT siblings. (I-J) Imaging of blood vessels in the posterior trunk using the transgenic reporter *flia*:GFP shows no difference in blood vessel morphology between WT siblings (I) and *pde* mutants (J). DA, Dorsal Aorta; PCV, Posterior Cardinal Vein; Se, Segmental Vessels.

To test the possible role of *ednraa* in pigment development, we used morpholino-mediated knockdown in wild-type embryos using the previously validated splice-blocking *ednraa*-morpholino (*ednraa*-MO1; Table 1 and Fig. 3B; [21]). As controls, we used injection of a random-sequence morpholino (cMO; standard control provided by Genetools) and the *ednrab* morpholino (*ednrab*-MO; Table 2; [21]), neither of which resulted in ectopic pigment cells, nor any other disruption to the early larval pigment pattern (Fig. 3C). In contrast, after injection of *ednraa*-MO1 we saw ectopic melanocytes and iridophores in the ventral medial pathway in a *pde*-like phenotype (Fig. 3D). As a further test, we designed a new translation blocking morpholino against *ednraa* (*ednraa*-MO2, Table 1); injection of this morpholino phenocopied the *pde* mutant pigment phenotype in a dose-dependent manner, but also was prone to non-specific deformations, likely due to off-target effects. Together, these data strongly support the hypothesis that the *pde* mutants disrupt *ednraa* function.

**Table 2.**
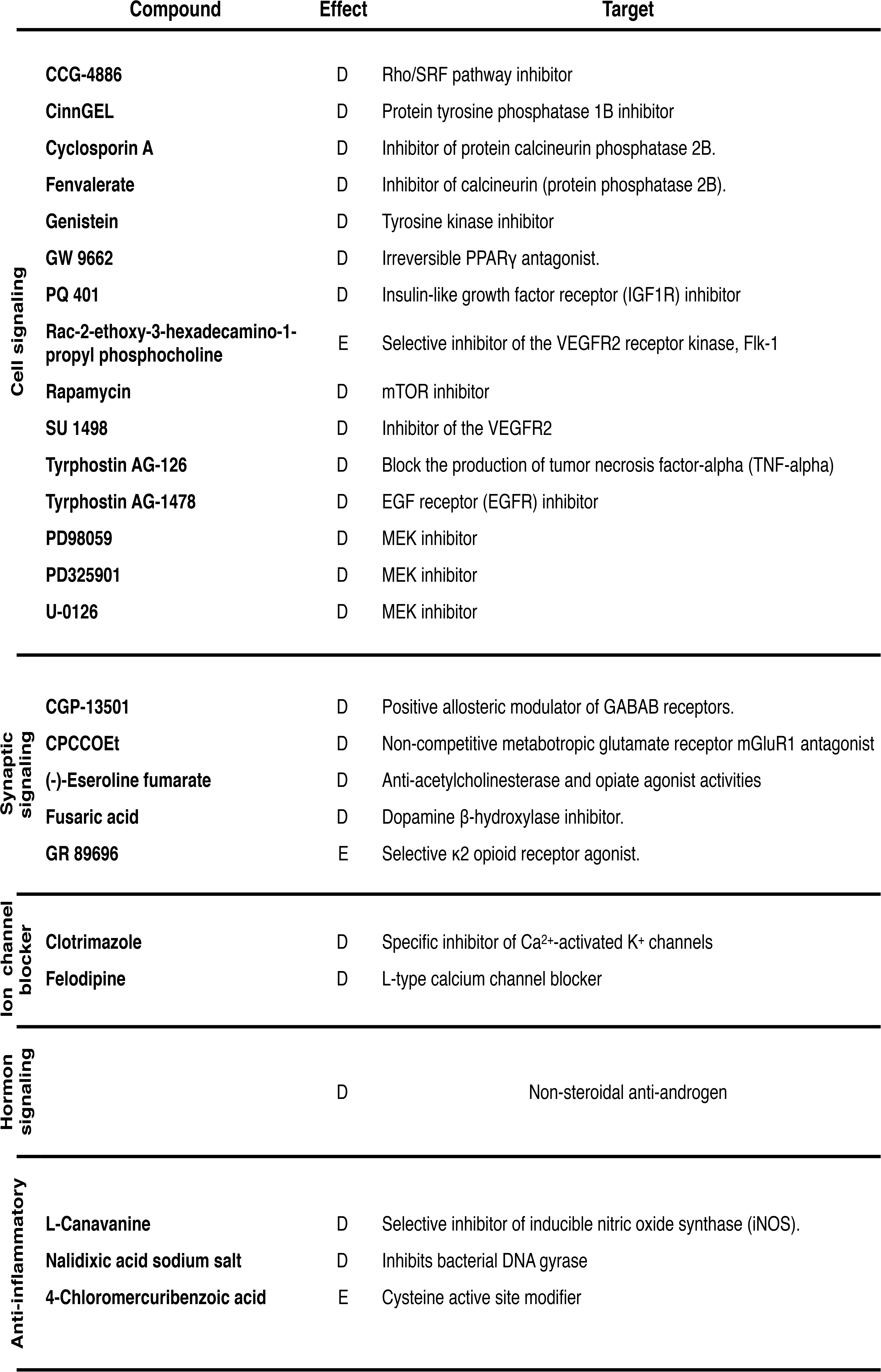
Screening of small molecules in *pde* mutants. Name and known targets of small molecules that enhance (E) or rescue (R) the *pde* phenotype.

To assess directly the link between *pde* mutations and disruption of *ednraa* function we amplified *ednraa* cDNAs from *pde* mutant alleles. In addition to the two original alleles, *pde^tj262^* and *pde^tv212^*, we also analysed *pde^hu4140^*, a third mutation identified in an independent screen at the Hubrecht Institute which showed a phenotype indistinguishable from the original alleles, and which failed to complement those original alleles (Supp. Fig. S2). This cDNA sequencing showed that *pde^hu4140^* has a single base transition mutation in the 3’ region of exon 5 (bp 847; AGA > TGA; Fig. 3B), which is predicted to result in a premature translation stop in Ednraa. The *pde^tj262^* and *pde^tv212^* alleles each showed deletions, of 103 bp and 135 bp respectively. cDNA sequence alignment indicates a deletion of exon 7 and a couple of extra bases, predicted to cause a frame shift of translation in exon 8 in *pde^tj262^*, while *pde^tv212^* has a deletion of exon 6 and a couple of extra bases which results in a frame shift of translation in exon 7 and 8 of Ednraa (Fig. 3B). Given that the *pde^tj262^* and *pde^tv212^* alleles were isolated from an ENU-mutagenesis screen and hence are likely to result from induced point mutations, we propose that the mutations in *pde^tj262^* and *pde^tv212^* are likely to affect key bases involved in *ednraa* splicing.

We used an *in situ* hybridization time-course between 6 and 72 hpf to assess the domain of *ednraa* expression, and in particular, whether it was detectable in neural crest or pigment cells. Our data was fully consistent with the earlier demonstration of *ednraa* expression in the developing blood vessels, but we saw no evidence of neural crest expression (Fig. 3E-H). We then tested whether blood vessels morphology was affected in *pde* mutants, which we will refer to as *ednraa* mutants from now on. One possible explanation for the pigment cells might be that blood vessel morphology might be disrupted in *ednraa* mutants, resulting in misplaced pigment cells. However, neither *in situ* hybridisation for *ednraa* (Fig. 3E-H), nor examination of trunk blood vessel morphology using the transgenic line Tg(*flia*:GFP (Fig. 3I,J) showed differences between WT and *ednraa* mutant embryos. We conclude that gross morphology of blood vessels is not affected in the *ednraa* mutants, but that the supernumerary and ectopic pigment cells in the *ednraa* mutants result from a noncell autonomous effect of endothelin signalling in the blood vessels.

### The *ednraa* phenotype does not result from neural crest cell transdifferentiation in the ventral trunk

The ventral medial pathway corresponds with the location of the nascent sympathetic ganglia, which form on the medial neural crest migration pathway in the vicinity of the dorsal aorta. We considered a transdifferentiation model in which neural crest cells fated to form sympathetic neurons switch to generating pigment cells, predicting that sympathetic neuron numbers would be reduced in *ednraa* mutants. However, immuno-detection of the early neuronal marker Elav1 (Hu neuronal RNA-binding protein) showed no differences in the number of sympathetic neurons between phenotypically WT embryos and their *ednraa* mutant siblings (Fig. 4A, 4B and 4K). Furthermore, we also tested whether other neural crest-derived neurons are affected in the trunk, but neither DRG sensory neuron nor enteric neuron numbers differed between *ednraa* mutants and their WT siblings. Thus, trunk DRGs contained around 3 neurons per ganglion (Fig. 4G, H and N;), while the posterior gut contained around 125 enteric neurons (Fig. 4I, J and O).

**Figure 4.**
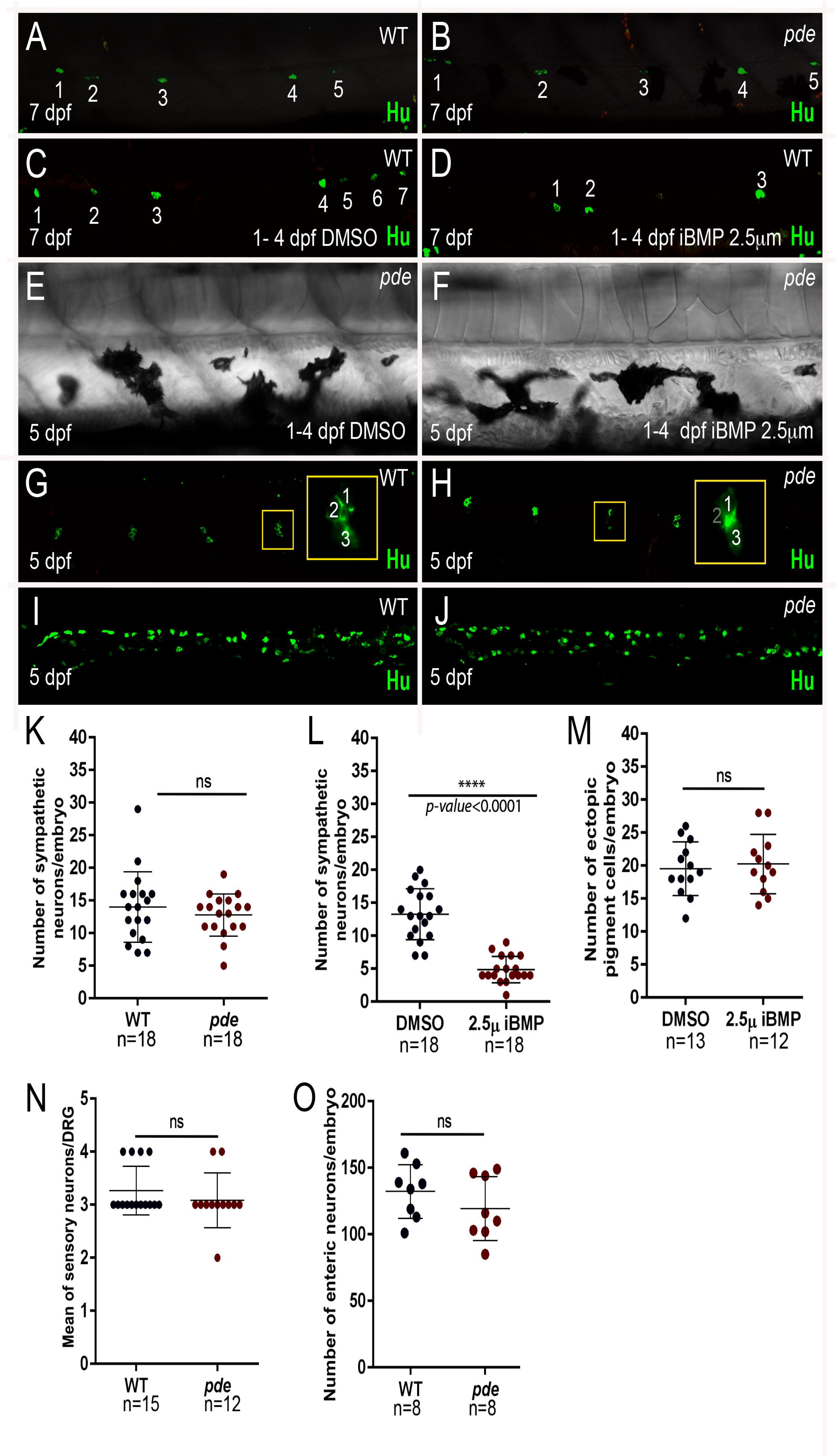
Neural crest-derived peripheral neurons are not reduced in *pde* mutants, but a role for BMP signaling in sympathetic neuron specification is conserved in zebrafish. Immunodetection of the neuronal marker Hu in 7 dpf WT embryos (**A**) and *pde* mutant siblings (**B**), shows no significant difference (ns) in the number of sympathetic neurons (K; WT mean±s.e.=14.0±1.27, n=18 and *pde*=12.78±0.76, n=18, n=18; p>0.05, two-tailed t-test). (**C-F**) Chemical inhibition of BMP signalling with dorsomorphin (iBMP 2.5 μM; 1-4 dpf treatment). Treatment of WT embryos shows a 63.25% reduction in the number of sympathetic neurons in comparison with 1% DMSO-treated controls (C, D and L; DMSO =13.28±0.91, n=18 and dorsomorphin=4.88±0.47, n=18; p<0.0001, two-tailed t-test). Quantification of the number of ectopic pigment cells in *pde* mutants treated with DMSO (E) or dorsomorphin (iBMP 2.5 μM; 1-4 dpf; F) shows no significant (ns) difference (M; DMSO mean±s.e.=19.54±1.12, n=13 and *pde* =20.25±1.30, n=12; p>0.05, two-tailed t-test). (**G-J**) Immunodetection of the neuronal marker Hu in 5 dpf WT embryos (G and I) and *pde* mutant siblings (H and J) shows no significant (ns) difference in the number of sensory neurons per dorsal root ganglion (DRG; G and H; WT=3.26±0.11, n=15 and *pde*=3.08±0.14, n=12; >0.05, two-tailed t-test) nor in enteric neurons in the posterior gut (I, J and O; WT =132.3±7.14, n=8 and *pde* =119.4±8.49, n=8; p>0.05, two-tailed t-test).

As a further test of the sympathetic neuron transfating hypothesis, we reasoned that if a key signal driving sympathetic neuron specification was reduced, this might result in enhanced numbers of ectopic pigment cells. In mammals and birds, secreted BMP signals from the dorsal aorta have been shown to induce sympathetic neurons [22–24], but it is not known if this mechanism is conserved in zebrafish. To test this, we treated zebrafish embryos with Dorsomorphin (2.5 μM), a well characterised BMP signalling inhibitor[25]. Given the well-known roles for BMP signaling in early patterning in the embryo, we chose a treatment window from 1-4 days post fertilisation (dpf). Although this treatment left the larvae looking morphologically normal, immunofluorescent detection of Elav1 showed that treated larvae had a strong reduction the number of sympathetic neurons compared to DMSO carrier-treated controls (Fig. 4C, D and L). Having shown that zebrafish sympathetic neurons were BMP-dependent, we then asked whether ectopic pigment cells in *ednraa* mutants were increased if sympathetic neuron specification was inhibited; using the same treatment conditions, we saw no enhancement of ectopic pigment cells in *ednraa* mutants treated with dorsomorphin compared to DMSO-treated controls (Fig. 4E, F and M). Thus, although we provide the first evidence to our knowledge that specification of zebrafish sympathetic neurons is BMP-dependent, we discount the hypothesis that the ectopic pigment cells in *ednraa* mutants result from transfating of sympathetic neurons.

### The *ednraa* phenotype results from localised increased neural crest cell proliferation in the ventral trunk, in the vicinity of the medial blood vessels

In order to identify the earliest stage at which ectopic pigment cells appear in the ventral trunk of *ednraa* mutants, we performed *in situ* hybridization with the melanocyte marker dct and the iridoblast marker ltk. Comparison of gene expression between WT and *ednraa* mutant embryos at 24, 30 and 35 hpf showed that ectopic/supernumerary expression of both *dct* and *ltk* is detected in the ventral trunk from 35 hpf (Fig. 5A-D).

**Figure 5.**
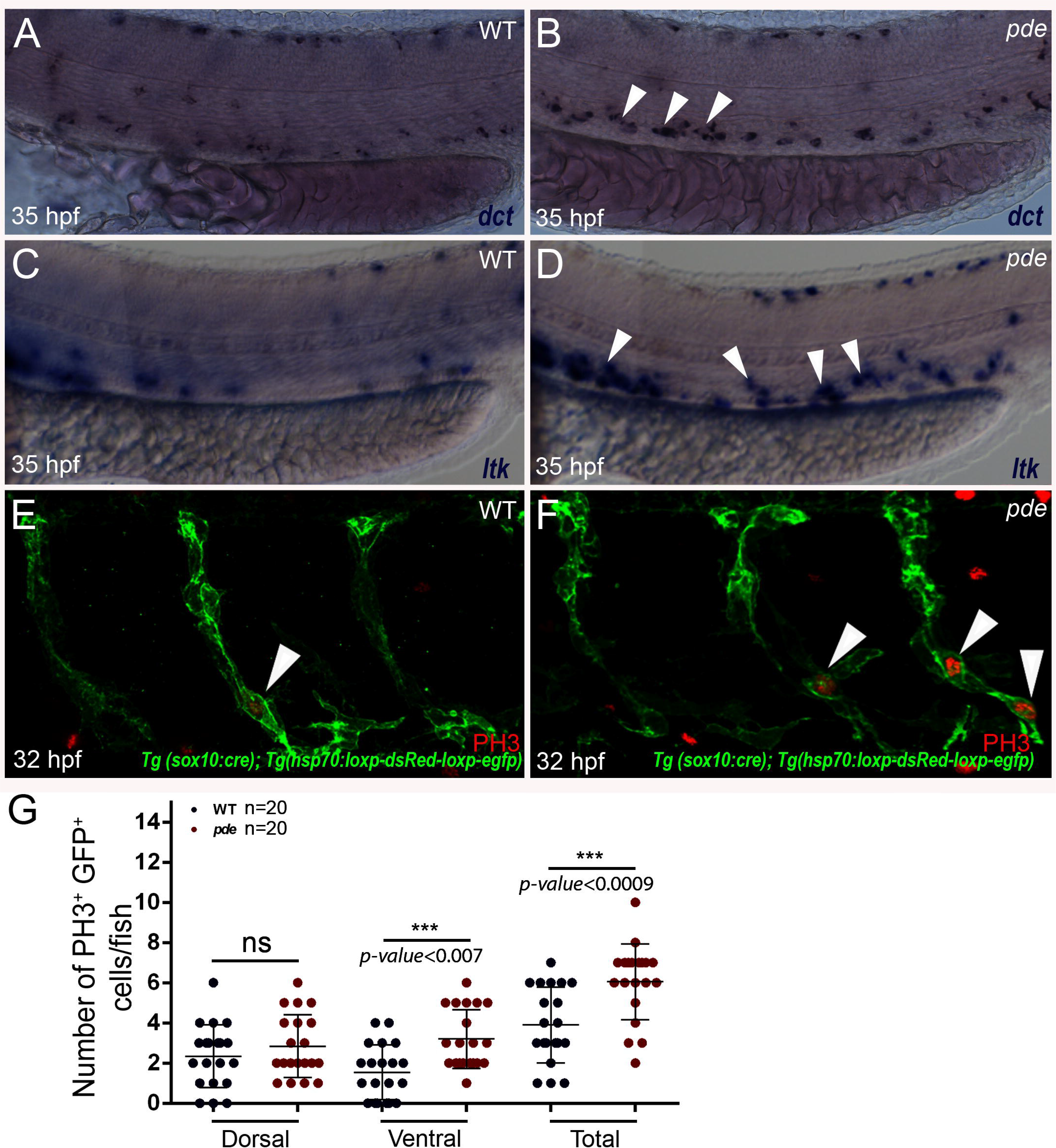
Ectopic pigment cells in *pde* mutants are detectable by 35 hpf, and generated by localised increased proliferation of neural crest-derived cells. (**A-D**) Whole mount in situ hybridization of 35 hpf WT (A and C) and *pde* mutant (B and D) embryos shows ectopic expression in *pde* mutants (white arrowheads in D) of melanocyte marker *dct* (A and B) and the iridophore marker *ltk* (C and D). (**E-G**) Immunodetection of the proliferation marker phosphohistone 3 (PH3) in neural crest derived cells (labelled with GFP due to *Tg(-4725sox10:cre)ba74; Tg(hsp:loxp-dsRed-loxp-LYN-EGFP))* of 32 hpf WT (E; white arrowhead) and *pde* mutant (F, white arrowheads) sibling embryos. Quantification of double positive GFP^+^ PH3^+^ cells in medial migratory pathway, shows a significant increase in *pde* mutants compared to WT siblings (Total; WT mean±s.e.=3.9±0.42, n=20 and *pde*=6.05±0.41, n=20; p<0.0009, two-tailed t-test). Subdividing this quantification of GFP^+^ PH3^+^ cells in the medial migratory pathway into those dorsal and ventral to the notochord shows that this increase is not detected on the dorsal medial pathway (dorsal double positive cells, WT=2.35±0.35, n=20 and *pde* =2.85±0.35, n=20; p<0.3188, two-tailed t-test), but is significantly increased on the ventral medial pathway WT=1.55±0.30, n=20 and *pde*=3.2±0.32, n=20; p<0.0007, two-tailed t-test).

Having defined the timing of appearance of ectopic cells in the ventral trunk of *ednraa* mutants, we tested whether this correlated with an increased proliferation of NC-derived cells in *ednraa* mutants. We used our *Tg(sox10:cre)* driver [26]combined with a *Tg(hsp70:loxP-dsRed-loxp-egfp)* red-green switch reporter [27] to label all neural crest cells with membrane tagged GFP, in conjunction with expression of the proliferation marker phosphohistone H3 by immunofluorescent labelling. Double labelling at 32 hpf, showed a 55% overall increase in NC-derived proliferating cells in *ednraa* mutants compared with WT siblings (Fig. 5E, F and G). Furthermore, over-proliferation of neural crest cells was detectable only in the ventral region of the medial pathway (Fig. 5G). Our results show that ectopic pigment cells in *ednraa* mutants result from overproliferation of NC-derived cells in a localised region of the medial pathway in the vicinity of the dorsal aorta, coinciding with the region of *ednraa* expression.

During the proliferation assays, we noted that the ventral proliferation of neural crest-derived cells was strongly clustered at the ventral end of the migrating streams (arrowheads, Fig. 5F). Furthermore, this association was true also of the ectopic pigment cells themselves, as shown by imaging of all NC-derived cells using the transgenic line *Tg(-4725sox10:cre)ba74; Tg(hsp:loxp-dsRed-loxp-EGFP)*(Supp. Fig. 3). This imaging confirmed the widespread ventral migration of neural crest-derived cells in *ednraa* mutants, comparable to that in WT siblings, at 35 hpf (Supp. Fig. 3A,B), reinforcing the conclusion from our WISH studies (Fig. 1 and 5). At 96 hpf, this imaging readily showed the ectopic pigment cells, but clearly revealed that their location is ventral to the notochord (and thus in the vicinity of the dorsal aorta), and also near the ventral projections of the spinal nerves (Supp. Fig. 3C,D).

### The formation of ectopic pigment cells in *ednraa* mutants requires ErbB signalling.

In order to identify possible mechanisms involved in the formation of the *ednraa* phenotype, we performed a small molecule screen of 1396 compounds using three different libraries: 1) the Sigma LOPAC library, which contains 1280 small organic ligands including marketed drugs and pharmaceutically relevant structures; 2) the Screen-WellTM Kinase Inhibitor Library that comprises 80 known kinase inhibitors; and 3) the Screen-WellTM Phosphatase Inhibitor Library, containing 33 phosphatase inhibitors of well-characterised activity. Screening was performed using the methodology of our previous screen for pigmentation modifiers [28]; we took advantage of the adult viability of the *ednraa* mutants, to perform the screen on *ednraa* mutant embryos. Thus *ednraa* mutant embryos were treated from 4 hpf to 96 hpf at a standard concentration of 10 μM and rescue or enhancement of *ednraa* mutant phenotype was systematically assessed at 4 dpf. After rescreening, we identified 23 compounds able to rescue the *ednraa* mutant phenotype, as well as 3 that enhance the phenotype (Table 2). Of these hits, four (Tyrphostin AG 1478, U0126, PD 98059 and PD325901) target the MAPK/ERK pathway: Tyrphostin AG-1478 is an inhibitor of the Epidermal Growth Factor Receptor (EGFR)[29] while U0126, PD98059 and PD325901 are highly selective inhibitors of MEK1/2 signalling [30–32]. Our previous work has shown that MEK inhibitors interfere with production of regenerative melanocytes, whilst not affecting direct developing pigment cells forming the early larval pattern in zebrafish [33, 34]. Treatment of *ednraa* mutants with each of two of the latter compounds shows a clear dose-response in the degree of rescue of the *ednraa* phenotype (Supp. Fig. 4).

The identification of Tyrphostin AG-1478 in our screen is especially notable, because it and the more specific EGFR inhibitor, PD158780, have been shown to affect APSC biology, but *not* embryonic pigment cells (except a small population contributing to the Lateral Stripe), in zebrafish [10]. Similarly, mutants for the epidermal growth factor receptor (EGFR)-like tyrosine kinase *erbb3b* (*picasso*), despite developing a normal embryonic/larval pigment pattern, fail to develop a normal adult pigment pattern and are unable to regenerate melanocytes after embryonic melanocyte ablation due to a lack of APSCs [10]. Thus inhibition of Erb signalling by either AG-1478 or PD158780 selectively affects the biology of NC-derived APSCs that give rise to adult and regenerative melanocytes. The rescue of the *ednraa* mutant by treatment with the Erb inhibitors suggested the exciting hypothesis that the ectopic cells in the *ednraa* mutants might result, not from embryonic pigment cells, but by precocious differentiation of APSCs. Interestingly, this role for ErbB signaling shows a tightly constrained time window, since inhibition of ErbB signalling with AG-1478 and PD158780 during window 9-48 hpf of embryonic development is sufficient for these effects [10]. Moreover, after melanocyte ablation throughout 24-72 hpf, melanocyte regeneration normally occurs by 5 dpf, but this is prevented when embryos are treated with AG-1478 during a 9-30 hpf window [7].

Thus, our hypothesis makes the testable prediction that rescue of the ectopic pigment cells in *ednraa* mutants would share the very specific temporal window known to regulate APSCs development. We treated *ednraa* mutant embryos with a range of concentrations (0.1 - 2.0 *μ* M) of PD158780 from 12-48 hpf (Fig. 6A-D). Quantification of the number of ectopic pigment cells showed that *ednraa* mutant embryos treated with PD158780 have a significant dose dependant-reduction in the number of ectopic cells compared to DMSO treated embryos (Fig. 6M). Moreover, shorter treatment with PD158780 revealed that the *ednraa* phenotype is effectively rescued when treatment is restricted to a 19-30 hpf window (Fig. 6E-H and 7M,), showing a striking match to the critical period for establishment of APSCs, while treatment after this window (24-30 hpf) does not rescue the *ednraa* phenotype (Fig. 6I-M). This data strongly supports the hypothesis that the source of ectopic pigment cells in *ednraa* mutants is likely to be APSCs and not embryonic pigment cells.

**Figure 6.**
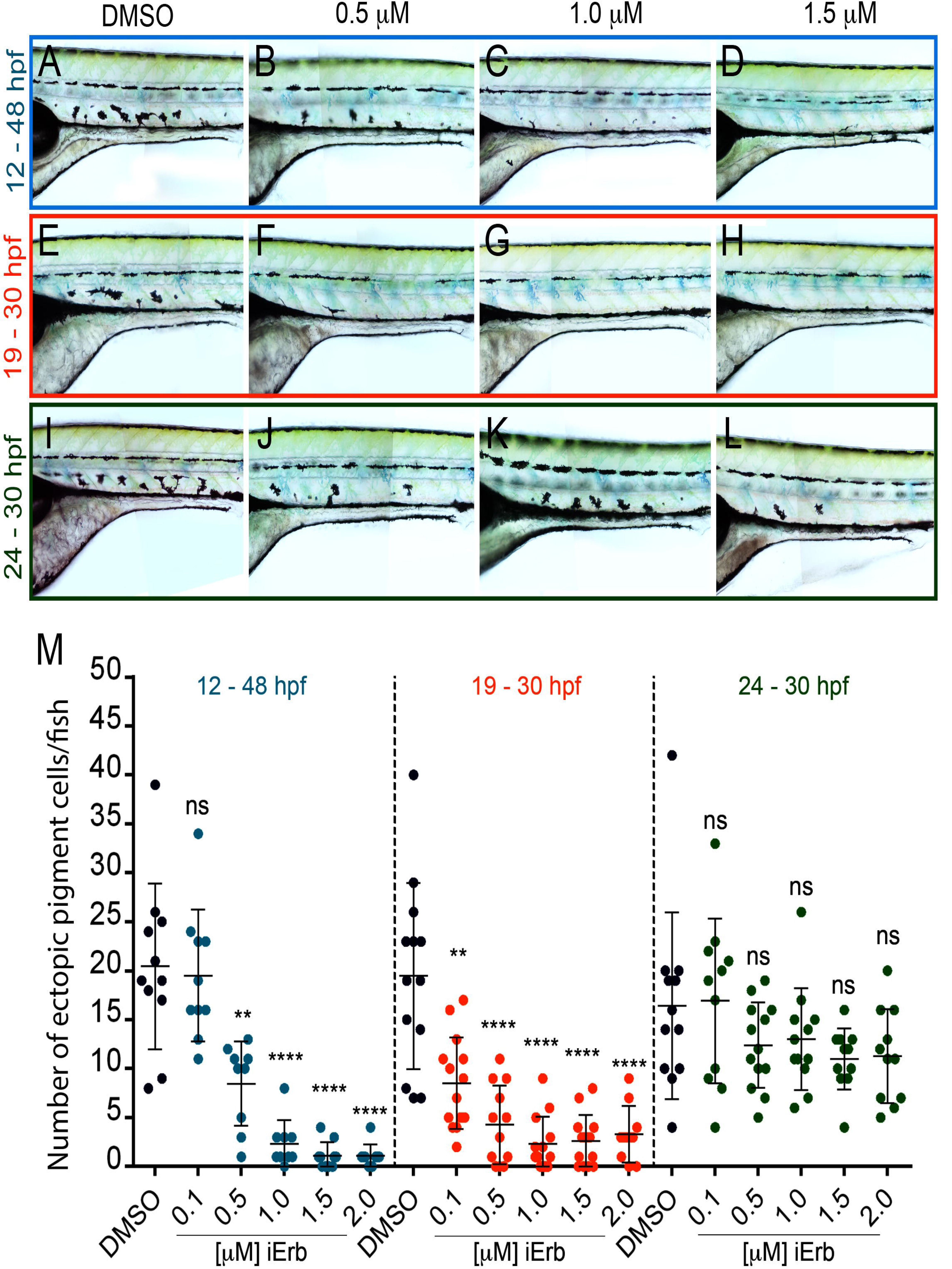
Chemical inhibition of Erb signalling rescues the pde phenotype. (**A-H**) Treatment of *pde* embryos with increasing concentrations of Erb inhibitor PD158780 (iErb; 0.5- 2.0 μM) or DMSO carrier control from 12-48 hpf (A-D), 19-30 hpf (E-H) and 24-30 hpf (I-L) hpf. Quantification of the number of ectopic pigment cells in the ventral medial pathway showed a decrease in the number of ectopic cells when embryos were treated from 12-48 hpf or just from 18-30 hpf, but not in a later 24-30 hpf time-window (M).

## Discussion

In this study we show that *ednraa* encodes one of two zebrafish Ednra orthologues, Ednraa. In mammals the EdnrA receptor binds selectively to Edn1 and Edn2, mediates vasoconstriction, and is overexpressed in many cancers[35]). In contrast, loss of function in mouse results in homeotic transformation of the lower jaw towards an upper jaw morphology, and in humans underlies Auriculocondylar syndrome (ACS [MIM 602483 and 614669])[36–39]. These studies did not identify pigmentation phenotypes, although an *Ednra* lacZ knockin mouse strain shows prominent expression in the hair follicles [36]. In contrast, *Ednrb* and *Edn3* mutants, as well as mutations in the Edn-processing enzyme Ece1, lack neural crest-derived melanocytes [40–42].

Analysis of the zebrafish genome identifies eleven components of endothelin signalling system: Four ligands, Edn1, Edn2, Edn3a and Edn3b; three Endothelin Converting Enzymes, Ece1, Ece2a, Ece2b that activate the ligands; and four receptors, Ednraa, Ednrab, Ednrba and Ednrbb [43]. In adult zebrafish, *ednrba* and *ece2b* loss-of-function mutants have all been shown to display reduced iridophores and broken stripes, indicating their coordinated role in iridophore development and pigment patterning, but no effect on embryonic pigment pattern [44, 45]. In contrast, *edn1* mutants and *ednraa* and *ednrab* morphants revealed disruption of the lower jaw, similar to the mammalian role in dorsoventral patterning, but did not examine pigmentation [21].

Here we identify a novel function for Ednraa signaling in pigment cell development. In silico translation and structural predictions for the *ednraa* alleles (Supp. Fig. 5A) indicate that the N-terminus and the early transmembrane domains might be intact, but that the other transmembrane domains and the C-terminus are absent. Our three *ednraa* mutant alleles show indistinguishable phenotypes, consistent with the similar predicted molecular effects of the mutations, and strongly indicating that the receptor is not functional. Consequently, we propose that these alleles are all likely null mutants.

Our experimental studies assess in turn a series of hypotheses regarding the embryonic basis of the ectopic pigment cell phenotype, initially exploring disrupted biology of embryonic (direct developing) pigment cells, before coming to the conclusion that the phenotype must result from disruption of APSC biology. The *ednraa* mutant phenotype is restricted to supernumerary and ectopic pigment cells in the ventral trunk and anterior tail; this spatial localization had been perplexing, but identification of the gene as *ednraa*, which is consistently expressed strongly in developing blood vessels (but not in NC) from well before the onset of detectable ectopic cells, helps to explain the restricted phenotype of *ednraa* mutants. We find no evidence that neural crest migration is disrupted, since not only is blood vessel morphology normal, but neural crest cell migration and patterning is normal except for the ectopic pigment cells themselves, and pigment cells are abundant in the Ventral and Yolk Sac Stripes. Similarly, we disprove the ‘mixed-fate’ hypothesis by showing using TEM that the ectopic cells are indistinguishable from those melanocytes and iridophores in the Yolk Sac Stripe where these two cell-types are consistently found in tight apposition. Instead, we conclude that the close association of the two cell-types reflects the natural tendency for iridophores to adhere to melanocytes (as seen in all locations in the early larval pattern). Finally, we find no evidence that the ectopic cells derive from transfating of sympathetic neurons, since neurons are unaffected in *ednraa* mutants and even when sympathetic neuron specification is inhibited using a BMP inhibitor, ectopic pigment cell number is unchanged.

However, these studies were unable to address the possible role of glial or progenitor cells in the ventral trunk. We reasoned that the supernumerary and ectopic cells were likely to be associated with increased proliferation of a subset of neural crest cells. Strikingly, we found that enhanced proliferation of neural crest did characterise the *ednraa* mutants, but that this proliferation was specifically associated with neural crest cells in ventral regions i.e. near the dorsal aorta, at around the time (35 hpf) that ectopic pigment cells begin to be identifiable. This is before the sympathetic ganglia show detectable differentiation and suggested that a subset of NC-derived cells undergoes ectopic or precocious proliferation. A key insight into the identity of these cells came from the observation that both ErbB inhibitors rescued the homozygous phenotype. Inhibition of Erb signaling *within a defined embryonic time window* by either AG-1478 or PD158780 selectively affects the biology of NC-derived APSCs that give rise to adult and regenerative melanocytes [7, 10]. We show that *ednraa* mutants are rescued by ErbB inhibitors in a dose- and time-dependent manner, and with a time-window overlapping that known to regulate APSC development. We conclude that the ectopic pigment cells in *ednraa* mutants result from precocious differentiation of pigment cells from APSCs, cells that would normally be quiescent until metamorphosis.

APSCs give rise to the numerous melanocyte and iridophores of the adult skin [11]. Our model would be consistent with the lack of ectopic xanthophores, since the evidence to date is that the majority of adult xanthophores derive from embryonic xanthophores and not from APSCs, although a small contribution from the stem cells is indicated by clonal analyses [11, 46]. Finally, the localised increase in neural crest cell proliferation that we document is also consistent with the activation of otherwise quiescent stem cells. APSCs in zebrafish have been closely-linked to the developing peripheral nervous system. One intriguing aspect of our data is that this proliferation is exclusively localised to the ventral trunk, consistent with the idea that the blood vessels form a key aspect of the stem cell niche, but surprising in that to date APSCs have been exclusively associated with the DRGs [9, 47]. We note that homozygous *ednraa* mutants are adult viable, but have no visible skin pigment pattern defect. Our data are consistent with a second source of APSCs, associated with the peripheral nervous system in the ventral trunk, likely the sympathetic ganglia. We propose a model in which these APSCs reside in a novel niche in close proximity to the medial blood vessels, which provide signals holding them in a quiescent state. In the *ednraa* mutants, these signals are reduced or absent, such that the APSCs become precociously activated. They then undergo proliferation and differentiate as melanocytes and iridophores. Many of these move into the Ventral Stripe location, but the excess cannot be accommodated in this stripe and remain ectopically located in the ventral trunk near the sympathetic ganglia (Fig. 7).

**Figure 7.**
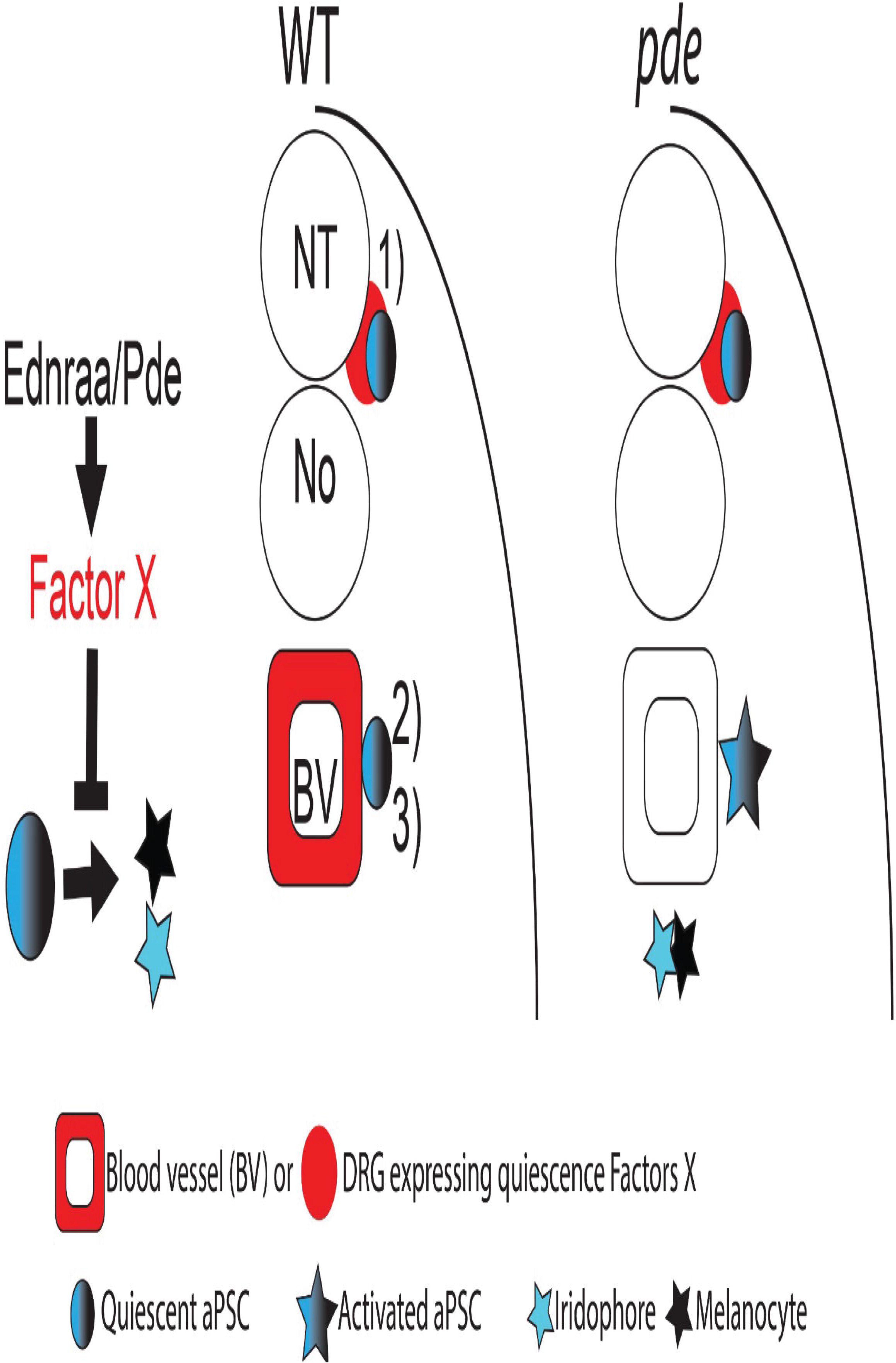
A second source of APSCs is held in a quiescent state by Ednraa-dependent factors from the blood vessels. Figure shows a model integrating our observations with current knowledge. 1) Dorsal root ganglia associated APSCs (APSC) are maintained in a quiescent state by local factors (red); 2) we propose a second source of APSCs in the vicinity of the medial blood vessels. Ednraa/pde activity in the blood vessels results in signals that hold this novel population in a quiescent state. In the *pde* mutants, these factors (red) are lost locally from the blood vessels and the APSCs become precociously activated, generating melanocytes and iridophores in their vicinity.

Although novel in the context of APSCs, blood vessels form an important part of adult stem cell niches in other contexts, especially that of adult Neural Stem Cells (NSCs) [48]. Indeed in the case of NSCs in the subependymal zone of mouse, blood vessel-mediated signals play a role in maintaining quiescence[48], although other sources for these quiescence signals have also been identified in the neurepithelial component of this niche [49–52]. In these studies of NSCs, notch signaling has been shown to be important in maintenance of quiescence, so it will be fascinating to compare the molecular signals derived from the vasculature that regulate APSC quiescence in the zebrafish. Our discovery of a second niche for APSCs and identification of the key role for blood vessels in controlling their behaviour provides an entry point for uncovering an important but currently understudied aspect of zebrafish pigment pattern formation.

Our work may also have implications for human disease. For example, Phakomatosis pigmentovascularis (PPV) is a rare mosaic disorder defined as the simultaneous occurrence of a widespread vascular nevus and an extensive pigmentary nevus, and associated with activating mutations of Gα subunits of heterotrimeric G proteins [53]. It is currently unknown if there is a common progenitor that gains an oncogenic mutation and leads to both large nevi and vascular proliferations, or if changes in one cell type impact upon another. Our study provides evidence that melanocytes can expand/differentiate in association with a modified blood vessel niche, making it conceivable that, for instance, oncogenic activation in blood vessels might result in non-cell autonomous activation of melanocyte stem cells in the niche to generate a nevus.

## Acknowledgments

We gratefully acknowledge funding support that enabled this research, specifically University of Bath Studentships (SC and JM), CONACyT grant 329640/384511 (KCS), and BBSRC grant BB/L00769X/1 and MRC grant MR/J001457/1 (RNK), and MRC Human Genetics Unit Programme (MC_PC_U127585840), European Research Council (ZF-MEL-CHEMBIO-648489) and L’Oreal-Melanoma Research Alliance (401181)(EEP). We thank Kerstin Howe and Mario Caccamo (Wellcome Trust Genome Campus) for their expert assistance in interpreting the zebrafish genome during the mapping, and Stefan Hans and Michael Brand for providing the Tg(hsp70:loxP-dsRed-loxp-egfp) red-green switch reporter.

## Author Contributions

**Supplementary figure S1. Melanocytes and iridophores in the WT yolk sac stripe are consistently separated from each other by double membranes**. Transmission electron photomicrographs of melanocytes and iridophores in the WT yolk sac stripe ectopic pigment cells in *pde* mutants. A and B show two examples of melanosomes (m) and reflecting. platelets (p) separated by a double membrane (white arrowheads).

**Supplementary figure S2. Complementation assay of the pde alleles**. Overview of early larval pigment phenotype at 5 dpf of *pde^tj262/tj262^* (A), *pde^tjj262/tv212^* and *pde^tj262/hu4140^*. All three allele combinations show ectopic melanocyte and iridophores in the ventral medial pathway of the posterior trunk.

**Supplementary figure S3. Migration of neural crest cells through the medial migratory pathway**. Labelling of neural crest derivatives with GFP using the transgenic line *Tg(-4725sox10:cre)ba74; Tg(hsp:loxp-dsRed-loxp-LYN-EGFP)* shows no difference between 35 hpf WT fish (A) and *pde* mutants (B), neural crest cells migrate ventrally in a intersegmental arrangement (white line in A and B). 96 hpf *pde* mutant larvae show ectopic pigment cells (white arrowheads) associated with the spinal nerve projections (white lines in C and D) that emerge from the dorsal root ganglion (red lines in C and D).

**Supplementary figure S4. Inhibition of MEK rescues the *pde* phenotype**. Treatment with increasing concentrations of the MEK inhibitors U0126 (2.5-7.6 μM) and PD 325901 (0.25 −0.75μM), from 6 - 96 hpf, shows increasing rescue of the ectopic pigment cells.

**Supplementary figure S5. In-silico translation and structural prediction for the *pde* alleles**. Schematic representation of the WT *ednraa* gene. Solid black line corresponds to intronic regions, but these are not shown to scale. Exons are numbered and shown in boxes. 5’ and 3’ UTRs are shown in orange, while coding exons are shown in alternating blue and green colour for visualisation purpose. Single letter a.a. sequence is shown. From top to bottom, the amino acid sequence provided by NCBI, and deduced sequences from in-silico translation of our sequenced cDNAs from AB, *pde^tj262^, pde^tv212^* and *pde^hu4140^* alleles. Solid green and blue lines link the corresponding exon to theamino acid sequence. Transmembrane domains (TD) are shown in dashed purple boxes and the extracellular domain (ED) and intracellular domains (ID) are labelled accordingly.

## References

1. Schartl M, Larue L, Goda M, Bosenberg MW, Hashimoto H, Kelsh RN. What is a vertebrate pigment cell? Pigment Cell Melanoma Res. 2016;29(1):8–14. doi: 10.1111/pcmr.12409. PubMed PMID: 26247887.

2. Irion U, Singh AP, Nusslein-Volhard C. The Developmental Genetics of Vertebrate Color Pattern Formation: Lessons from Zebrafish. Curr Top Dev Biol. 2016;117:141–69. doi: 10.1016/bs.ctdb.2015.12.012. PubMed PMID: 26969976.

3. Kelsh RN. Genetics and evolution of pigment patterns in fish. Pigment Cell Res. 2004;17(4):326–36. PubMed PMID: 15250934.

4. Kelsh RN, Harris ML, Colanesi S, Erickson CA. Stripes and belly-spots – a review of pigment cell morphogenesis in vertebrates. Semin Cell Dev Biol. 2009;20(1):90–104. Epub 2008/11/04. doi: 10.1016/j.semcdb.2008.10.001S1084-9521 (08)00096-7 [pii]. PubMed PMID: 18977309; PubMed Central PMCID: PMC2744437.

5. Parichy DM. Animal pigment pattern: an integrative model system for studying the development, evolution, and regeneration of form. Semin Cell Dev Biol. 2009;20(1):63–4. Epub 2009/01/17. doi: S1084-9521(08)00155-9 [pii] 10.1016/j.semcdb.2008.12.010. PubMed PMID: 19146966.

6. Singh AP, Nusslein-Volhard C. Zebrafish stripes as a model for vertebrate colour pattern formation. Curr Biol. 2015;25(2):R81–92. doi: 10.1016/j.cub.2014.11.013. PubMed PMID: 25602311.

7. Hultman KA, Budi EH, Teasley DC, Gottlieb AY, Parichy DM, Johnson SL. Defects in ErbB-dependent establishment of adult melanocyte stem cells reveal independent origins for embryonic and regeneration melanocytes. PLoS Genet. 2009;5(7):e1000544. Epub 2009/07/07. doi: 10.1371/journal.pgen.1000544. PubMed PMID: 19578401; PubMed Central PMCID: PMC2699538.

8. Johnson SL, Nguyen AN, Lister JA. mitfa is required at multiple stages of melanocyte differentiation but not to establish the melanocyte stem cell. Dev Biol. 2011;350(2):405–13. Epub 2010/12/15. doi: S0012-1606(10)01245-5 [pii] 10.1016/j.ydbio.2010.12.004. PubMed PMID: 21146516; PubMed Central PMCID: PMC3040983.

9. Dooley CM, Mongera A, Walderich B, Nusslein-Volhard C. On the embryonic origin of adult melanophores: the role of ErbB and Kit signalling in establishing melanophore stem cells in zebrafish. Development. 2013;140(5):1003–13. Epub 2013/02/01. doi: 10.1242/dev.087007 dev.087007 [pii]. PubMed PMID: 23364329.

10. Budi EH, Patterson LB, Parichy DM. Embryonic requirements for ErbB signaling in neural crest development and adult pigment pattern formation. Development. 2008;135(15):2603–14. Epub 2008/05/30. doi: dev.019299 [pii] 10.1242/dev.019299. PubMed PMID: 18508863.

11. Singh AP, Dinwiddie A, Mahalwar P, Schach U, Linker C, Irion U, et al. Pigment Cell Progenitors in Zebrafish Remain Multipotent through Metamorphosis. Dev Cell. 2016. doi: 10.1016/j.devcel.2016.06.020. PubMed PMID: 27453500.

12. Svetic V, Hollway GE, Elworthy S, Chipperfield TR, Davison C, Adams RJ, et al. Sdf1a patterns zebrafish melanophores and links the somite and melanophore pattern defects in choker mutants. Development. 2007;134(5):1011–22. PubMed PMID: 17267445.

13. Raible DW, Wood A, Hodsdon W, Henion PD, Weston JA, Eisen JS. Segregation and early dispersal of neural crest cells in the embryonic zebrafish. Dev Dyn. 1992;195(1):29–42.

14. Kelsh RN, Schmid B, Eisen JS. Genetic analysis of melanophore development in zebrafish embryos. Dev Biol. 2000;225(2):277–93. PubMed PMID: 10985850.

15. Lister JA, Robertson CP, Lepage T, Johnson SL, Raible DW. nacre encodes a zebrafish microphthalmia-related protein that regulates neural-crest-derived pigment cell fate. Development. 1999;126(17):3757–67. PubMed PMID: 10433906.

16. Lopes SS, Yang X, Muller J, Carney TJ, McAdow AR, Rauch GJ, et al. Leukocyte tyrosine kinase functions in pigment cell development. PLoS Genet. 2008;4(3):e1000026. Epub 2008/03/29. doi: 10.1371/journal.pgen.1000026. PubMed PMID: 18369445.

17. Minchin JE, Hughes SM. Sequential actions of Pax3 and Pax7 drive xanthophore development in zebrafish neural crest. Dev Biol. 2008;317(2):508–22. Epub 2008/04/18. doi: S0012-1606(08)00177-2 [pii] 10.1016/j.ydbio.2008.02.058. PubMed PMID: 18417109.

18. Kelsh RN, Brand M, Jiang YJ, Heisenberg CP, Lin S, Haffter P, et al. Zebrafish pigmentation mutations and the processes of neural crest development. Development. 1996;123:369–89.

19. Petratou K, Camargo-Sosa K, Al Jabri R, Nagao Y, Kelsh RN. Transcript and protein detection methodologies for neural crest research on whole mount zebrafish and medaka. In: Schwarz Q, Wiszniak S, editors. Methods relevant to Neural Crest Cell research. Methods in Molecular Biology Springer; in press.

20. Geisler R, Rauch GJ, Geiger-Rudolph S, Albrecht A, van Bebber F, Berger A, et al. Large-scale mapping of mutations affecting zebrafish development. BMC Genomics. 2007;8:11. Epub 2007/01/11. doi: 1471-2164-8-11 [pii] 10.1186/1471-2164-8-11. PubMed PMID: 17212827; PubMed Central PMCID: PMC1781435.

21. Nair S, Li W, Cornell R, Schilling TF. Requirements for Endothelin type-A receptors and Endothelin-1 signaling in the facial ectoderm for the patterning of skeletogenic neural crest cells in zebrafish. Development. 2007;134(2):335–45. doi: 10.1242/dev.02704. PubMed PMID: 17166927.

22. Reissmann E, Ernsberger U, Francis-West P, Rueger D, Brickell P, Rohrer H. Involvement of bone morphogenetic protein-4 and bone morphogenetic protein-7 in the differentiation of the adrenergic phenotype in developing sympathetic neurons. Development. 1996; 122(7):2079–88.

23. Schneider C, Wicht H, Enderich J, Wegner M, Rohrer H. Bone morphogenetic proteins are required in vivo for the generation of sympathetic neurons. Neuron. 1999;24(4):861–70.

24. Saito D, Takahashi Y. Sympatho-adrenal morphogenesis regulated by the dorsal aorta. Mech Dev. 2015;138 Pt 1:2–7. doi: 10.1016/j.mod.2015.07.011. PubMed PMID: 26235279.

25. Yu PB, Hong CC, Sachidanandan C, Babitt JL, Deng DY, Hoyng SA, et al. Dorsomorphin inhibits BMP signals required for embryogenesis and iron metabolism. Nat Chem Biol. 2008;4(1):33–41. Epub 2007/11/21. doi: nchembio.2007.54 [pii] 10.1038/nchembio.2007.54. PubMed PMID: 18026094; PubMed Central PMCID: PMC2727650.

26. Rodrigues FS, Doughton G, Yang B, Kelsh RN. A novel transgenic line using the Cre-lox system to allow permanent lineage-labeling of the zebrafish neural crest. Genesis. 2012;50(10):750–7. Epub 2012/04/24. doi: 10.1002/dvg.22033. PubMed PMID: 22522888.

27. Hans S, Freudenreich D, Geffarth M, Kaslin J, Machate A, Brand M. Generation of a non-leaky heat shock-inducible Cre line for conditional Cre/lox strategies in zebrafish. Dev Dyn. 2011;240(1):108–15. Epub 2010/12/01. doi: 10.1002/dvdy.22497. PubMed PMID: 21117149.

28. Colanesi S, Taylor KL, Temperley ND, Lundegaard PR, Liu D, North TE, et al. Small molecule screening identifies targetable zebrafish pigmentation pathways. Pigment Cell Melanoma Res. 2012;25(2):131–43. Epub 2012/01/19. doi: 10.1111/j.1755-148X.2012.00977.x. PubMed PMID: 22252091.

29. Levitzki A, Gazit A. Tyrosine kinase inhibition: an approach to drug development. Science. 1995;267(5205):1782–8. PubMed PMID: 7892601.

30. Barrett SD, Bridges AJ, Dudley DT, Saltiel AR, Fergus JH, Flamme CM, et al. The discovery of the benzhydroxamate MEK inhibitors CI-1040 and PD 0325901. Bioorg Med Chem Lett. 2008;18(24):6501–4. doi: 10.1016/j.bmcl.2008.10.054. PubMed PMID: 18952427.

31. Dudley DT, Pang L, Decker SJ, Bridges AJ, Saltiel AR. A synthetic inhibitor of the mitogen-activated protein kinase cascade. Proc Natl Acad Sci U S A. 1995;92(17):7686–9. PubMed PMID: 7644477; PubMed Central PMCID: PMCPMC41210.

32. Favata MF, Horiuchi KY, Manos EJ, Daulerio AJ, Stradley DA, Feeser WS, et al. Identification of a novel inhibitor of mitogen-activated protein kinase kinase. J Biol Chem. 1998;273(29):18623–32. PubMed PMID: 9660836.

33. Anastasaki C, Estep AL, Marais R, Rauen KA, Patton EE. Kinase-activating and kinase-impaired cardio-facio-cutaneous syndrome alleles have activity during zebrafish development and are sensitive to small molecule inhibitors. Hum Mol Genet. 2009;18(14):2543–54. Epub 2009/04/21. doi: ddp186 [pii] 10.1093/hmg/ddp186. PubMed PMID: 19376813; PubMed Central PMCID: PMC2701326.

34. Grzmil M, Whiting D, Maule J, Anastasaki C, Amatruda JF, Kelsh RN, et al. The INT6 Cancer Gene and MEK Signaling Pathways Converge during Zebrafish Development. PLoS ONE. 2007;2(9):e959. Epub 2007/09/27. doi: 10.1371/journal.pone.0000959. PubMed PMID: 17895999.

35. Cong N, Li Z, Shao W, Li J, Yu S. Activation of ETA Receptor by Endothelin-1 Induces Hepatocellular Carcinoma Cell Migration and Invasion via ERK1/2 and AKT Signaling Pathways. J Membr Biol. 2016;249(1-2):119–28. doi: 10.1007/s00232-015-9854-1. PubMed PMID: 26501871.

36. Gordon CT, Petit F, Kroisel PM, Jakobsen L, Zechi-Ceide RM, Oufadem M, et al. Mutations in endothelin 1 cause recessive auriculocondylar syndrome and dominant isolated question-mark ears. Am J Hum Genet. 2013;93(6):1118–25. doi: 10.1016/j.ajhg.2013.10.023. PubMed PMID: 24268655; PubMed Central PMCID: PMCPMC3853412.

37. Miller CT, Schilling TF, Lee K, Parker J, Kimmel CB. sucker encodes a zebrafish Endothelin-1 required for ventral pharyngeal arch development. Development. 2000;127(17):3815–28. PubMed PMID: 10934026.

38. Sato T, Kurihara Y, Asai R, Kawamura Y, Tonami K, Uchijima Y, et al. An endothelin-1 switch specifies maxillomandibular identity. Proc Natl Acad Sci US A. 2008; 105(48): 18806–11. doi: 10.1073/pnas.0807345105. PubMed PMID: 19017795; PubMed Central PMCID: PMCPMC2596216.

39. Clouthier DE, Garcia E, Schilling TF. Regulation of facial morphogenesis by endothelin signaling: insights from mice and fish. Am J Med Genet A. 2010;152A(12):2962–73. doi: 10.1002/ajmg.a.33568. PubMed PMID: 20684004; PubMed Central PMCID: PMCPMC2974943.

40. Baynash AG, Hosoda K, Giaid A, Richardson JA, Emoto N, Hammer RE, et al. Interaction of endothelin-3 with endothelin-B receptor is essential for development of epidermal melanocytes and enteric neurons. Cell. 1994;79(7):1277–85.

41. Hosoda K, Hammer RE, Richardson JA, Baynash AG, Cheung JC, Giaid A, et al. Targeted and natural (piebald-lethal) mutations of endothelin-B receptor gene produce megacolon associated with spotted coat color in mice. Cell. 1994;79(7): 1267–76.

42. Yanagisawa H, Yanagisawa M, Kapur RP, Richardson JA, Williams SC, Clouthier DE, et al. Dual genetic pathways of endothelin-mediated intercellular signaling revealed by targeted disruption of endothelin converting enzyme-1 gene. Development. 1998;125(5):825–36. PubMed PMID: 98119793.

43. Braasch I, Volff JN, Schartl M. The endothelin system: evolution of vertebrate-specific ligand-receptor interactions by three rounds of genome duplication. Mol Biol Evol. 2009;26(4):783–99. Epub 2009/01/29. doi: msp015 [pii] 10.1093/molbev/msp015. PubMed PMID: 19174480.

44. Krauss J, Astrinidis P, Frohnhofer HG, Walderich B, Nusslein-Volhard C. transparent, a gene affecting stripe formation in Zebrafish, encodes the mitochondrial protein Mpv17 that is required for iridophore survival. Biology open. 2013;2(7):703–10. doi: 10.1242/bio.20135132. PubMed PMID: 23862018; PubMed Central PMCID: PMCPMC3711038.

45. Parichy DM, Mellgren EM, Rawls JF, Lopes SS, Kelsh RN, Johnson SL. Mutational analysis of endothelin receptor b1 (rose) during neural crest and pigment pattern development in the zebrafish Danio rerio. Dev Biol. 2000;227(2):294–306. doi: 10.1006/dbio.2000.9899. PubMed PMID: 11071756.

46. McMenamin SK, Bain EJ, McCann AE, Patterson LB, Eom DS, Waller ZP, et al. Thyroid hormone-dependent adult pigment cell lineage and pattern in zebrafish. Science. 2014;345(6202): 1358–61. doi: 10.1126/science.1256251. PubMed PMID: 25170046; PubMed Central PMCID: PMCPMC4211621.

47. Singh AP, Schach U, Nusslein-Volhard C. Proliferation, dispersal and patterned aggregation of iridophores in the skin prefigure striped colouration of zebrafish. Nat Cell Biol. 2014;16(6):607–14. Epub 2014/04/30. doi: 10.1038/ncb2955. PubMed PMID: 24776884.

48. Ottone C, Krusche B, Whitby A, Clements M, Quadrato G, Pitulescu ME, et al. Direct cell-cell contact with the vascular niche maintains quiescent neural stem cells. Nat Cell Biol. 2014; 16(11): 1045–56. doi: 10.1038/ncb3045. PubMed PMID: 25283993; PubMed Central PMCID: PMCPMC4298702.

49. Kawai H, Kawaguchi D, Kuebrich BD, Kitamoto T, Yamaguchi M, Gotoh Y, et al. Area-Specific Regulation of Quiescent Neural Stem Cells by Notch3 in the Adult Mouse Subependymal Zone. J Neurosci. 2017;37(49):11867–80. doi: 10.1523/JNEUROSCI.0001-17.2017. PubMed PMID: 29101245.

50. Engler A, Rolando C, Giachino C, Saotome I, Erni A, Brien C, et al. Notch2 Signaling Maintains NSC Quiescence in the Murine Ventricular-Subventricular Zone. Cell Rep. 2018;22(4):992–1002. doi: 10.1016/j.celrep.2017.12.094. PubMed PMID: 29386140.

51. Alunni A, Krecsmarik M, Bosco A, Galant S, Pan L, Moens CB, et al. Notch3 signaling gates cell cycle entry and limits neural stem cell amplification in the adult pallium. Development. 2013;140(16):3335–47. doi: 10.1242/dev.095018. PubMed PMID: 23863484; PubMed Central PMCID: PMCPMC3737716.

52. Chapouton P, Skupien P, Hesl B, Coolen M, Moore JC, Madelaine R, et al. Notch activity levels control the balance between quiescence and recruitment of adult neural stem cells. J Neurosci. 2010;30(23):7961–74. doi: 10.1523/JNEUROSCI.6170-09.2010. PubMed PMID: 20534844.

53. Thomas AC, Zeng Z, Riviere JB, O’Shaughnessy R, Al-Olabi L, St-Onge J, et al. Mosaic Activating Mutations in GNA11 and GNAQ Are Associated with Phakomatosis Pigmentovascularis and Extensive Dermal Melanocytosis. J Invest Dermatol. 2016;136(4):770–8. doi: 10.1016/j.jid.2015.11.027. PubMed PMID: 26778290; PubMed Central PMCID: PMCPMC4803466.

